# The Multidrug Resistance Transporter P-glycoprotein Confers Resistance to Ferroptosis Inducers

**DOI:** 10.1101/2023.02.23.529736

**Authors:** William J. E. Frye, Lyn M. Huff, José M. González Dalmasy, Paula Salazar, Rachel M. Carter, Ryan T. Gensler, Dominic Esposito, Robert W. Robey, Suresh V. Ambudkar, Michael M. Gottesman

## Abstract

Ferroptosis is a form of cell death caused by direct or indirect inhibition of glutathione peroxidase 4 that leads to lethal lipid peroxidation. Several small molecule ferroptosis inducers (FINs) have been reported, yet little information is available regarding resistance mechanisms, particularly their interaction with the ATP-binding cassette (ABC) transporters P-glycoprotein (P-gp, ABCB1) and ABCG2. Given the role that ABC transporters play in absorption, distribution, and excretion of many drugs, characterizing these interactions could provide information regarding oral bioavailability and brain penetration and may predict drug-drug interactions. Using ferroptosis-sensitive A673 cells transfected to express P-gp or ABCG2, we found that P-gp overexpression was able to confer resistance to FIN56 and the erastin derivatives imidazole ketone erastin and piperazine erastin. Results were confirmed with OVCAR8-derived NCI/ADR-RES cells that overexpress P-gp, where the P-gp inhibitor valspodar completely inhibited resistance to the FINs. P-gp-mediated resistance to imidazole ketone erastin and piperazine erastin was also reversed in UO-31 renal cancer cells by CRISPR-mediated knockout of *ABCB1*. At a concentration of 10 μM, the FINs ML-162, GPX inhibitor 26a, and PACMA31 were able to increase intracellular rhodamine 123 fluorescence over 10-fold in P-gp-expressing MDR-19 cells and GPX inhibitor 26a was able to increase intracellular purpurin-18 fluorescence over 4-fold in ABCG2-expressing R-5 cells. Expression of P-gp may reduce the efficacy of these FINs in cancers that express the transporter and may prevent access to sanctuary sites such as the brain. The ability of some FINs to inhibit P-gp and ABCG2 suggests potential drug-drug interactions.

**SIGNIFICANCE STATEMENT:** While several small-molecule ferroptosis inducers have been described, little work has addressed potential interactions with ABC transporters such as P-glycoprotein or ABCG2 that might limit bioavailability or brain penetration. We find that the ferroptosis inducers FIN56, imidazole ketone erastin, and piperazine erastin are substrates of P-glycoprotein. ML-162, GPX inhibitor 26a, and PACMA31 were found to inhibit P-glycoprotein, while GPX inhibitor 26a was additionally able to inhibit ABCG2, suggesting the potential for drug-drug interactions.

## INTRODUCTION

Ferroptosis is an iron-dependent form of non-apoptotic cell death arising from direct or indirect inhibition of glutathione peroxidase 4 (GPX4), leading to lipid peroxidation and unsustainable levels of reactive oxygen species (ROS)(Stockwell et al., 2017). One of the first reported inducers of ferroptosis, erastin, was described before the concept of ferroptosis was completely understood. Erastin was found to selectively kill transformed human foreskin fibroblasts expressing mutant HRAS compared to isogenic cells expressing wild-type HRAS (Dolma et al., 2003). The mechanism of cell death induced by erastin was not apoptosis, as the hallmarks of apoptotic cell death such as annexin V staining and caspase 3 cleavage were not observed, although death was accompanied by cell membrane permeabilization (Dolma et al., 2003). The term “ferroptosis” was later coined for this novel form of cell death as iron chelators and antioxidants were found to potentiate erastin-mediated toxicity, suggesting an iron-dependent increase in ROS was responsible (Dixon et al., 2012). Additionally, ferroptosis could not be inhibited by caspase inhibitors and was found to occur independently of the apoptosis effector proteins Bak and Bax (Dixon et al., 2012).

The target of erastin was identified to be the cystine/glutamate antiporter, system x^-^_c_ (Dixon et al., 2012). Inhibition of the antiporter leads to depletion of glutathione and inactivation of GPX4 (Yang et al., 2014). Subsequent to the discovery of erastin, other small molecules have been developed to induce ferroptosis by direct or indirect inhibition of GPX4. These ferroptosis inducers (FINs) include modified forms of erastin such as erastin2 (Dixon et al., 2014), imidazole ketone erastin (Zhang et al., 2019) and piperazine erastin (Yang et al., 2014), as well as the inhibitors FIN56 (Shimada et al., 2016), RSL3 (Yang and Stockwell, 2008), FINO2 (Gaschler et al., 2018), PACMA31(Chen et al., 2020), GPX4 inhibitor 26a (Xu et al., 2021), and ML-162 and ML-210 (Weïwer et al., 2012).

In an attempt to identify cancers that might be effectively treated by ferroptosis induction, Yang and colleagues tested erastin toxicity in 117 cancer cell lines and identified renal cell carcinomas as being particularly sensitive to ferroptosis (Yang et al., 2014). This intrigued us, as renal cell carcinomas are often positive for expression of P-glycoprotein (P-gp, encoded by the *ABCB1* gene), an ATP-binding cassette (ABC) multidrug efflux pump that confers drug resistance (Kanamaru et al., 1989; Gottesman et al., 2002). Some reports have also suggested that expression levels of another ABC transporter, ABCG2 (encoded by the *ABCB2* gene), also expressed in kidney cancers, can predict overall survival in patients with clear cell renal carcinoma (Wang et al., 2017). Both P-gp and ABCG2 localize to the gastrointestinal tract as well as to other barrier sites such as the blood-brain barrier (BBB) (Durmus et al., 2015). Expression at these sites is linked to their role in limiting the oral bioavailability of chemotherapy drugs and brain penetration of several targeted therapies (Vlaming et al., 2009; Durmus et al., 2015). We thus sought to characterize the interactions between small-molecule ferroptosis inducers and the transporters P-gp and ABCG2.

## MATERIALS AND METHODS

### Chemicals

Erastin, doxorubicin, rhodamine 123, and FIN56 were purchased from Sigma-Aldrich (St. Louis, MO). Erastin2, imidazole ketone erastin, RSL3, ML-162, FINO2, GPX4 inhibitor 26a, JKE-1674, and JKE-1716 were from Cayman Chemical (Ann Arbor, MI). Valspodar was obtained from MedChemExpress (Monmouth Junction, NJ). Romidepsin was from Selleck Chemicals (Houston, TX). Purpurin-18 was purchased from Frontier Scientific (Logan, UT). Piperazine erastin was from TargetMol (Wellesly Hills, MA). SN-38 was obtained from LKT Laboratories (St. Paul, MN). RSL3 was purchased from Tocris (Minneapolis, MN). Fumitremorgin C (FTC) was synthesized in-house by the Developmental Therapeutics Program at the National Institutes of Health (Bethesda, MD).

### Cell Lines

OVCAR8, NCI/ADR-RES and UO-31 cells were obtained from the Division of Cancer Treatment and Diagnosis Tumor Repository, National Cancer Institute (Frederick, MD) and are grown in RPMI-1640 with 10% FBS, glutamine and Pen/Strep. A673 cells (from ATCC, Manassas, VA) were seeded and transfected with empty vector (EV) or vector containing fulllength, human *ABCB1* or *ABCG2* using Lipofectamine 2000 (Invitrogen, Waltham, MA). Cells were selected with hygromycin, and clones were isolated by limiting dilution. Selected clones were grown in DMEM with 10% fetal calf serum, glutamine and Pen/Strep as well as 300 μg/ml hygromycin to maintain expression of the transporters. P-gp-overexpressing MDR-19 cells and ABCG2-overexpressing R-5 cells were derived from HEK293 cells and have been previously characterized and described (Robey et al., 2011). Transfected HEK293 cells were grown in MEM with 10% fetal calf serum, glutamine and Pen/Strep along with 2 mg/ml G418 to maintain expression of the transporters. All cell lines were routinely tested for mycoplasma using the MycoAlert PLUS Kit (Promega, Madison, WI) test kit and their identities were confirmed by STR analysis (performed by ATCC, Manassas, VA).

### Oligonucleotides

The following oligonucleotides (generated by Eurofins, Inc., Louisville, KY) were used in this study:

ABCB1-START: 5’-GGGGACAACTTTGTACAAAAAAGTTGGCACCATGGATCTTGAAGGGGACCGCAATGG
ABCB1-END: 5’-GGGGACAACTTTGTACAAGAAAGTTGATTATGCTAGCTGGCGCTTTGTTCCAGCCTGG
ABCG2-START: 5’-GGGGACAACTTTGTACAAAAAAGTTGGCACCATGTCTTCCAGTAATGTCGAAGTTTTTATCCC
ABCG2-END: 5’-GGGGACAACTTTGTACAAGAAAGTTGATTAAGAATACTTTTTAAGAAATAACAATTTCAG

### Generation of Entry Clones

Entry clones for *ABCB1* and *ABCG2* were constructed by PCR amplification of cDNA sequences flanked by Gateway Multisite recombination sites (Thermo Fisher Scientific, Waltham, MA). PCR was carried out with 200 nM of each oligo listed in the table above using Phusion polymerase (New England Biolabs, Ipswich, MA) under standard conditions and an extension time of 180 seconds. PCR products were cleaned using the QiaQuick PCR purification kit (Qiagen, Germantown, MD). The final PCR products were recombined into Gateway Donor vector pDonr-253 using the Gateway BP recombination reaction using the manufacturer’s protocols. The subsequent Entry clones were sequence verified throughout the entire cloned region.

### Subcloning for Mammalian Expression Constructs

Gateway Multisite LR recombination was used to construct the final mammalian expression constructs from the Entry clones using the manufacturer’s protocols (Thermo Fisher Scientific, Waltham, MA). The Gateway Destination vector used was pDest-305 (Addgene. Watertown, MA, #161895), a mammalian expression vector containing a Gateway attR4-attR2 cassette based on a modified version of pcDNA3.1. This vector backbone contains a hygromycin resistance marker for antibiotic selection. A human elongation factor 1 (EF1) promoter was introduced using a Gateway att4-att1 Entry clone (Addgene, Watertown, MA, #162920). Final expression clones were verified by restriction analysis and maxiprep DNA was prepared using the Qiaprep Maxiprep kit (Qiagen, Germantown, MD).

### Generation of ABCB1 Knockout UO-31 Cells

CRISPR-mediated knockout of *ABCB1* in UO-31 cells was achieved by co-transfecting cells with knockout and homology-directed repair vectors for *ABCB1* (obtained from Santa Cruz Biotechnology, Dallas, TX) using Lipofectamine 2000 (Invitrogen) and subsequent selection with puromycin (3 μg/ml). Knockout clones were subsequently isolated, and loss of P-gp was verified by flow cytometry.

### Cytotoxicity Assays

Cells were seeded in opaque white, 96-well plates at a density of 2500 cells/well and allowed to attach overnight. Cells were then treated with increasing concentrations of the desired compound and incubated for 72 h. The concentration at which 50% of cell growth was inhibited (GI50) was determined using Cell TiterGlo (Promega) according to the manufacturer’s instructions. Where noted, cytotoxicity assays were performed with 10 μM valspodar to inhibit P-gp.

### Flow Cytometry Assays

To measure cell surface expression of P-gp or ABCG2, trypsinized cells were incubated for 20 min. at room temperature in 2% bovine serum albumin/PBS with phycoerythrin-labeled UIC-2 antibody or phycoerythrin-labeled 5D3 antibody, respectively, according to the manufacturer’s instructions (both from ThermoFisher, Grand Island, NY). Cells were also incubated with the corresponding phycoerythrin-labeled isotype control—IgG2a kappa for P-gp and IgG2b kappa for ABCG2 (both from ThermoFisher). P-gp or ABCG2 transporter activity was measured using rhodamine 123 or purpurin-18, respectively (Robey et al., 2004). Cells were trypsinized and incubated for 30 min in complete medium (phenol red-free Richter’s medium with 10% FCS and penicillin/streptomycin) with the desired fluorescent substrate (0.5 μg/ml rhodamine 123 to detect P-gp or 15 μM purpurin-18 to detect ABCG2) in the presence or absence of 25 μM concentrations of the desired FIN or a positive control inhibitor (10 μM valspodar for P-gp or 10 μM fumitremorgin C for ABCG2) for 30 min at 37°C in 5% CO2. Subsequently, cells were washed and incubated in substrate-free medium for 1 h at 37°C continuing with or without inhibitor. Cells were subsequently analyzed with a FACSCanto flow cytometer (BD Biosciences, San Jose, CA) and data analysis was performed using FloJo v10.4.2 (FlowJo LLC, Ashland OR).

### ATPase Assay

The ATPase assay was performed as described previously (Ambudkar, 1998). Total membrane vesicles were prepared from High Five insect cells overexpressing P-gp. The membranes were diluted with ATPase assay buffer (50 mM MES-Tris, pH 6.8 containing 50 mM KCl, 5 mM NaN3, 1 mM EGTA, 10 mM MgCl2, 2 mM DTT and 1 mM oubain) to reach a final concentration of 100 μg/mL and were incubated with the compounds at the noted concentrations for 10 min in the presence or absence of 0.3 mM sodium orthovanadate. The addition of 5 mM ATP (5 mM) started the reaction (20 min at 37°C) after which SDS (2.5% final concentration) was added to terminate the reaction. The amount of inorganic phosphate released was quantified andthe results were reported as percentage of vanadate-sensitive ATPase activity with DMSO.

## RESULTS

### Generation and Characterization of A673 Cells That Overexpress P-gp or ABCG2

As the A673 cell line was reported to be sensitive to FINs (Yang et al., 2014) and does not express very high levels of P-gp or ABCG2, we transfected this cell line with either empty vector (A673 EV) or vectors containing the genes encoding human P-gp (A673 B1) or ABCG2 (A673 G2). We selected single clones with high levels of P-gp or ABCG2 based on measurement of antibody staining by flow cytometry and further characterized positive clones. As seen in Fig. 1A, A673 B1 or A673 G2 cells were found to have much higher levels of the transporter proteins, as shown by increased staining with UIC-2 antibody or 5D3 antibody (orange histogram), respectively, compared to A673 EV cells which were negative for both transporters. P-gp-overexpressing MDR-19 cells and ABCG2-overexpressing R-5 cells served as positive controls (data not shown). Functional assays were also used to confirm transporter activity. A673 B1 cells readily transported rhodamine 123 and A673 G2 cells demonstrated increased purpurin-18 efflux, as shown by the left shift of the blue histogram for the two substrates, compared to empty vector-transfected cells (A673 EV), as illustrated in Fig. 1B.

**Fig. 1.**
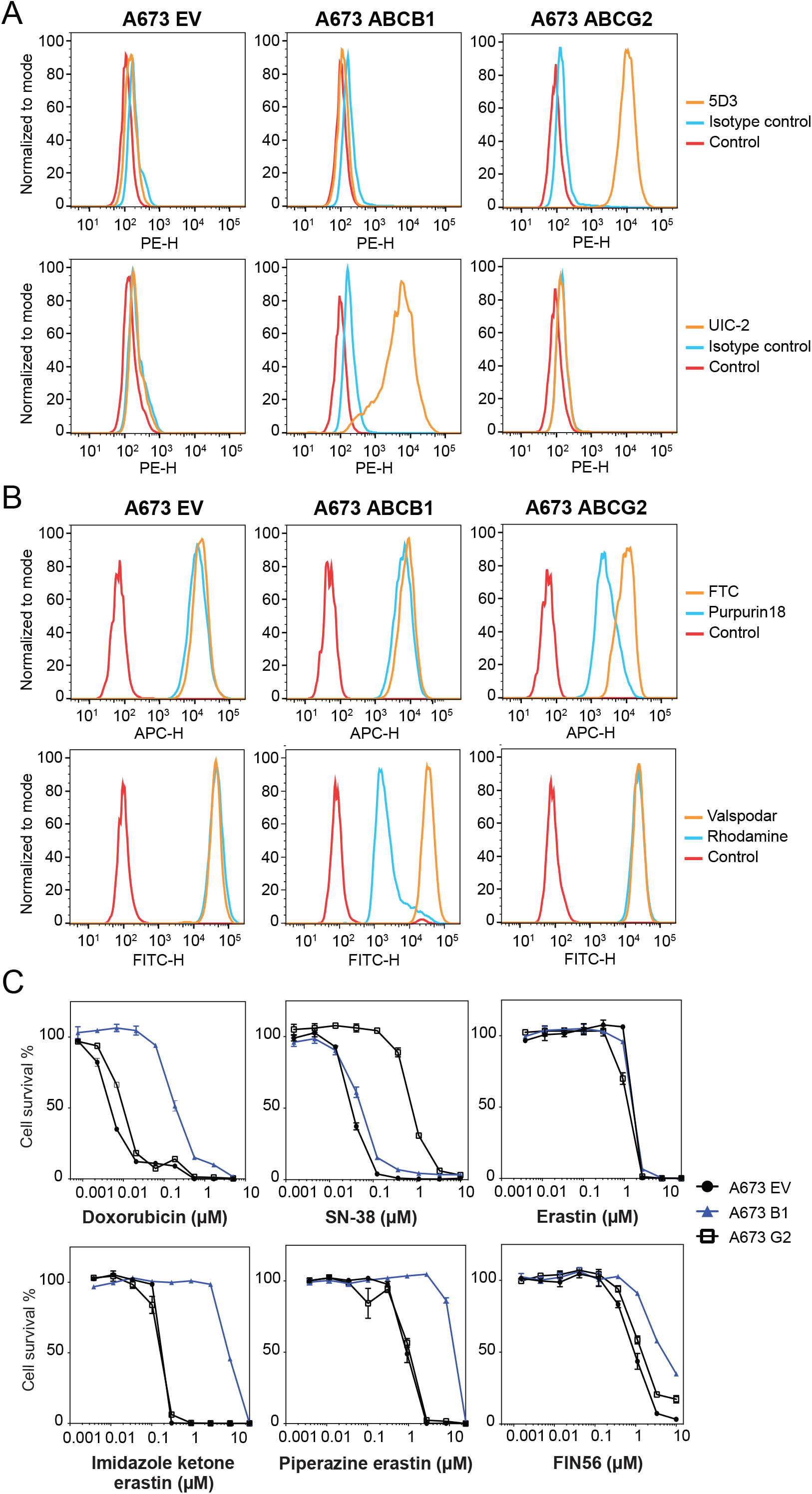
Characterization of A673 cells transfected to express P-gp or ABCG2. (A) Trypsinized A673 EV, B1 or G2 cells were incubated with 2% bovine serum albumin/PBS containing phycoerythrin-labeled antibody to detect ABCG2 (5D3) or ABCB1 (UIC-2), or the corresponding isotype control antibody for 20 min after which cells were washed in PBS. Control cells (no antibody) are denoted by red curves, isotype control staining is denoted by blue curves, and staining with specific transporter antibodies is denoted by orange curves (ABCG2 top row, P-gp bottom row). Results from one of three independent experiments are shown. (B) Trypsinized A673 EV, B1, and G2 cells were incubated with rhodamine 123 (0.5 μg/ml, for detection of P-gp) or purpurin-18 (15 μM, for detection of ABCG2) with or without appropriate inhibitor (10 μM valspodar for P-gp; 10 μM FTC for ABCG2) for 30 min after which media was removed and replaced with substrate-free medium continuing with or without inhibitor for an additional 1 h. Cell autofluorescence (control) is denoted by red histograms, substrate efflux is denoted by blue histograms and cells with substrate and inhibitor are denoted by orange histograms. Results from one of three independent experiments are shown. (C) Three-day cytotoxicity assays were performed on A673 EV, B1, and G2 cells with doxorubicin, SN-38, erastin, imidazole ketone erastin, piperazine erastin and FIN56. Results from one of three independent experiments are shown and results are summarized in Table 1.

To verify that transporter levels were adequate to confer resistance to known substrates, we performed cytotoxicity assays with the P-gp substrate doxorubicin as well as the ABCG2 substrate SN-38. As seen in Fig. 1C and Table 1, A673 B1 cells were resistant to doxorubicin, whereas A673 G2 cells exhibited little to no resistance to this compound. A673 G2 cells displayed increased resistance to SN-38. Having confirmed that the transfected cells expressed high levels of the desired transporters and that the transporters were indeed functional, we proceeded to use them to characterize the ability of P-gp and ABCG2 to confer resistance to FINs.

**Table 1.** Cross resistance profile of transfected A673 cells^*a*^

| **Compound (μM)** | **A673 EV GI<sub>50</sub>** | **A673 B1 GI<sub>50</sub>** | **RR** | **A673 G2 GI<sub>50</sub>** | **RR** |
| --- | --- | --- | --- | --- | --- |
| SN-38 | 0.0018 ± 0.001 | 0.0024 ± 0.0004 | 1.3 | 0.042 ± 0.00013 | 23 |
| Doxorubicin | 0.010 ± 0.004 | 0.24 ± 0.075 | 24 | 0.017 ± 0.005 | 1.7 |
| Erastin | 2.2 ± 0.42 | 3.4 ± 1.2 | 1.5 | 1.9 ± 0.78 | 0.86 |
| Erastin2 | 0.083 ± 0.004 | 0.17 ± 0.043 | 2 | 0.087 ± 0.024 | 1 |
| Imidazole ketone erastin | 0.31 ± 0.25 | 11.4 ± 5.4 | 38 | 0.33 ± 0.24 | 1.1 |
| Piperazine erastin | 1.6 ± 0.416 | 11 ± 2 | 6.9 | 1.4 ± 0.28 | 0.88 |
| FIN56 | 0.70 ± 0.43 | 3.9 ± 2.8 | 5.6 | 0.99 ± 0.65 | 1.4 |
| JKE-1674 | 1.0 ± 0.20 | 1.9 ± 0.18 | 1.9 | 2.7 ± 0.11 | 2.7 |
| RSL3 | 0.055 ± 0.010 | 0.12 ± 0.008 | 2.2 | 0.06 ± 0.003 | 1.1 |
| FinO<sub>2</sub> | 2.8 ± 1.7 | 2.8 ± 1.7 | 1 | 3.0 ± 1.6 | 1.1 |
| PACMA31 | 0.047 ± 0.056 | 0.07 ± 0.056 | 1.5 | 0.045 ± 0.039 | 0.96 |
| ML-210 | 0.16 ± 0.031 | 0.33 ± 0.028 | 2.1 | 0.20 ± 0.037 | 1.2 |
| GPX4 inhibitor 26a | 0.077 ± 0.036 | 0.14 ± 0.050 | 1.8 | 0.093 ± 0.020 | 1.2 |
| ML-162 | 0.095 ± 0.013 | 0.20 ± 0.057 | 2.1 | 0.12 ± 0.026 | 1.3 |
| JKE-1716 | 1.4 ± 0.38 | 2.5 ± 0.43 | 1.8 | 0.81 ± 0.054 | 0.58 |

<sup>a</sup>Results presented are mean GI<sub>50</sub> values +/- SEM. Relative resistance (RR) value was determined by dividing the GI<sub>50</sub> value for A673 cells expressing a transporter by the GI<sub>50</sub> value for the A673 EV cells. Three independent experiments were performed.

### P-gp Overexpression Confers Resistance to Modified Erastin Derivatives

A673 EV, A673 B1 and A673 G2 cells were then used in 3-day cytotoxicity assays to determine whether the transporters could confer resistance to the FINs. While resistance due to expression of ABCG2 or P-gp was not seen to most FINs examined, this was not true for some erastin derivatives that had been modified to improve water solubility. Overexpression of P-gp conferred relatively high levels of resistance to imidazole ketone erastin and piperazine erastin (Fig. 1C). FIN56 appeared to be a weak P-gp substrate, as A673 B1 cells were about 6-fold resistant (Table 1). ABCG2 overexpression did not confer appreciable resistance to any of the FINs examined.

Since P-gp overexpression appeared to confer resistance to some FINs, we validated the results in parental OVCAR8 ovarian cancer cells and P-gp-overexpressing NCI/ADR-RES cells that were derived from OVCAR8 cells by selection with doxorubicin. As shown in Fig. 2A, OVCAR8 cells do not express P-gp, as determined with the P-gp-specific monoclonal antibody UIC-2, while NCI/ADR-RES cells express high levels of the transporter, as shown by increased staining with the UIC-2 antibody in NCI/ADR-RES cells. In this model system, when we performed cytotoxicity assays in the presence or absence of 10 μM valspodar, a P-gp inhibitor, we observed that P-gp overexpression conferred resistance to imidazole ketone erastin, piperazine erastin, and FIN56, and that valspodar reversed the resistance. P-gp overexpression in the NCI/ADR-RES line was not found to confer resistance to erastin or erastin2 (Fig. 2B and Table S1), in agreement with the results from the A673 cells.

**Fig. 2.**
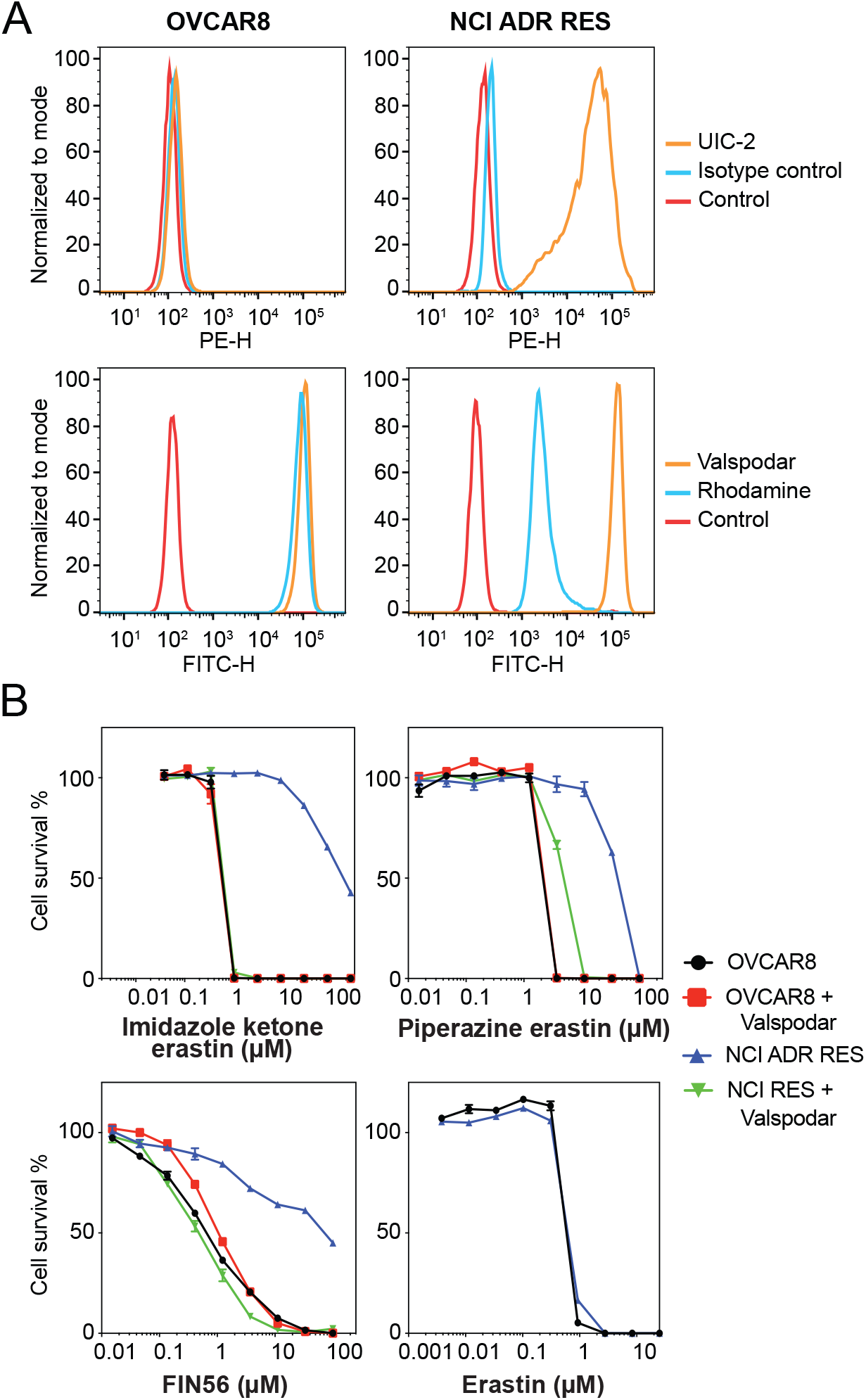
The OVCAR8 and NCI/ADR-RES cell line pair confirm P-gp substrates. (A) Trypsinized OVCAR8 or NCI/ADR-RES cells were incubated with 2% bovine serum albumin/PBS containing phycoerythrin-labeled UIC-2 antibody to isotype control antibody for 20 min after which cells were washed in PBS and read on a flow cytometer. Control cells (no antibody) are denoted by red curves, isotype control staining is denoted by blue curves, and P-gp staining is denoted by orange curves (ABCG2 top row, P-gp bottom row). Results from one of three independent experiments are shown. (B) Three-day cytotoxicity assays were performed on OVCAR8 and NCI/ADR-RES cells with imidazole ketone erastin, piperazine erastin, FIN56 or erastin. Where noted, the P-gp inhibitor valspodar was added at a concentration of 10 μM. Results from one of three independent experiments are shown and results are summarized in Table S1.

### Deletion of *ABCB1* in UO-31 Cells Increases Sensitivity to Erastin Derivatives

The renal carcinoma cell line UO-31 is known to have detectable levels of P-gp and displays rhodamine efflux (Lee et al., 1994). To determine if expression of P-gp in this cell line is high enough to confer resistance to imidazole ketone erastin or piperazine erastin, two of the best substrates for P-gp, we performed CRISPR-mediated deletion of *ABCB1* and selected two clones that had lost expression. As shown in Fig. 3A, while UO-31 cells do stain positively with the UIC-2 antibody, as shown by increased staining with the UIC-2 antibody (orange histogram), the two knockout clones, B11 and 1F4, no longer react with the antibody, as detected by flow cytometry. Additionally, we found that the knockout clones no longer efflux the P-gp substrate rhodamine 123, as shown by increased intracellular fluorescence of rhodamine 123 (blue histogram) in the clones, suggesting that the *ABCB1* gene had been deleted. The knockout clones also demonstrate increased sensitivity to the P-gp substrate romidepsin (Fig. 3B). When we performed cytotoxicity assays with imidazole ketone erastin and piperazine erastin, the knockout clones displayed increased sensitivity to both compounds by about 3-to 4-fold compared to parental cells; however, no difference in sensitivity to erastin was noted (Table S2). Thus, even relatively low levels of P-gp may cause resistance to the erastin analogs.

**Fig. 3.**
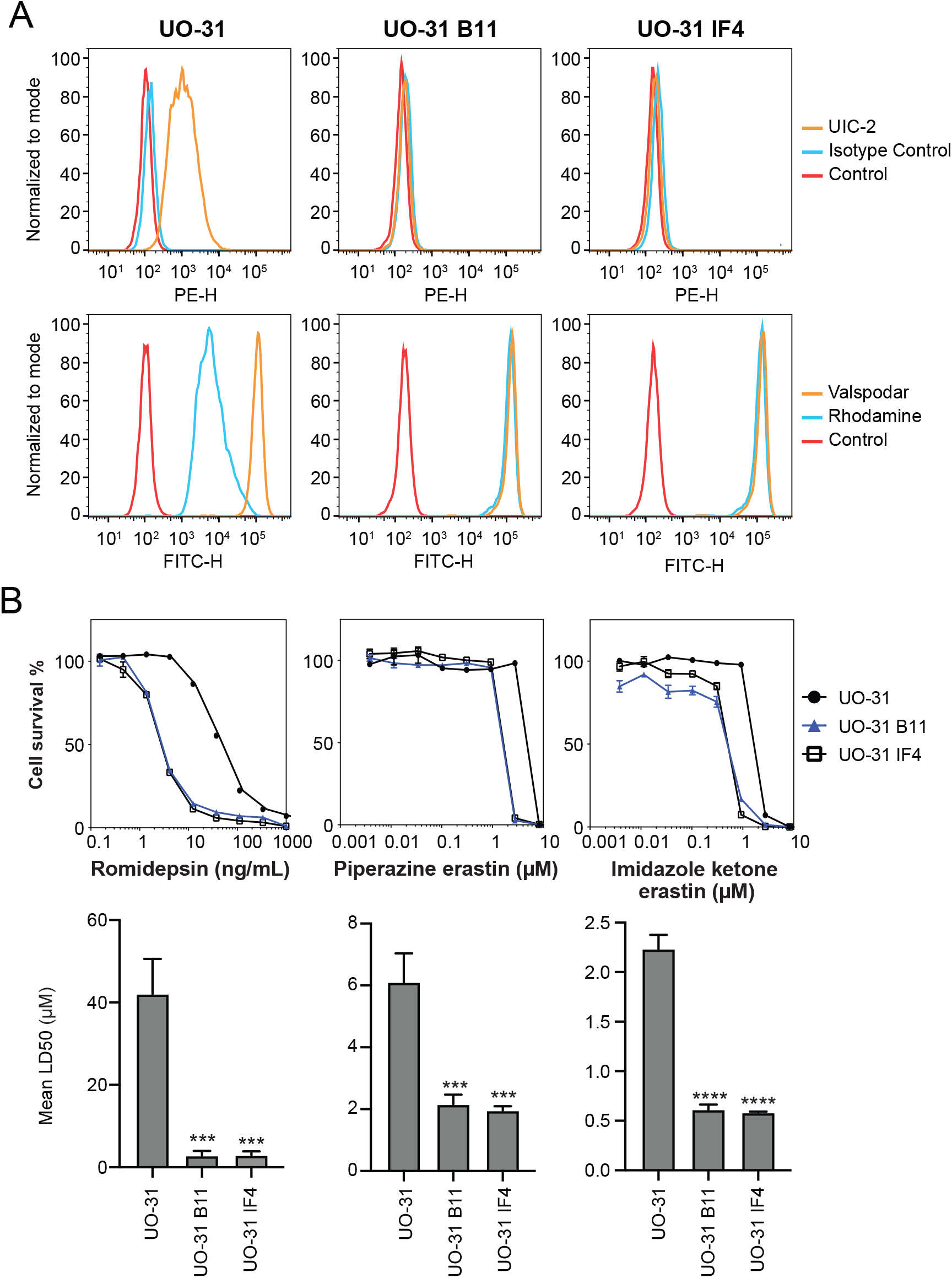
CRISPR-mediated deletion of ABCB1 sensitizes UO-31 cells to FINs. (A) Top row: UO-31 cells or the *ABCB1* knockout clones (B11, 1F4) were trypsinized and incubated with 2% bovine serum albumin/PBS containing phycoerythrin-labeled UIC-2 antibody or isotype control antibody for 20 min after which cells were washed in PBS and read on a flow cytometer. Control cells (no antibody) are denoted by red curves, isotype control staining is denoted by blue curves and staining with UIC-2 is denoted by orange curves. Bottom row: Cells were incubated with rhodamine 123 (0.5 μg/ml) with or without 10 μM valspodar for 30 min after which media was removed and replaced with substrate-free medium continuing with or without inhibitor for an additional 1 h. Cell autofluorescence (control) is denoted by red histograms, rhodamine efflux by blue histograms and cells with rhodamine and inhibitor are denoted by orange histograms. Results from one of three independent experiments are shown. (B) Three-day cytotoxicity assays were performed on UO-31 cells or the *ABCB1* knockout clones with rhodamine, imidazole ketone erastin or piperazine erastin. Results from one of 3 independent experiments are shown. GI50 values from 3 independent experiments are shown under the representative graphs. Significance was determined by a one-way ANOVA with a test for multiple comparisons. Asterisks denote significant difference from the parental UO-31 cell line, where p<0.001(***), or p<0.0001(****).

### Ferroptosis Inducers Stimulate the ATPase Activity of P-gp

The effect of erastin, imidazole ketone erastin, and piperazine erastin on the ATPase activity of P-gp was subsequently examined. While several P-gp substrates and some inhibitors have been shown to stimulate the ATPase activity of P-gp, not all substrates do so. We found that all three of the compounds stimulated the ATPase activity of P-gp to a degree comparable to that of verapamil, which is used as a positive control (Fig. 4). However, stimulation by erastin did not achieve statistical significance. The ATPase stimulation serves as a confirmation of the interaction between imidazole ketone erastin and piperazine erastin and P-gp, given the close connection between ATP hydrolysis and substrate efflux (Ambudkar et al., 1997).

**Fig. 4.**
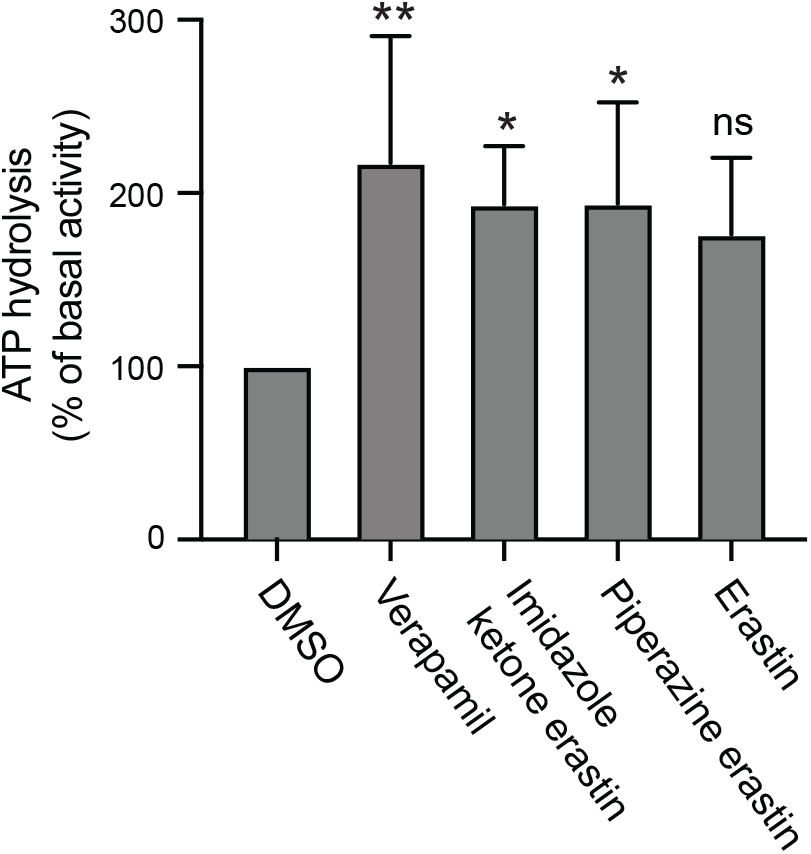
Effect of erastin derivatives on ATPase activity of P-gp. The effect of imidazole ketone erastin, piperazine erastin and erastin or the vanadate-sensitive ATPase activity of P-gp was determined as outlined in Materials and Methods. Basal ATPase activity was compared to that in the presence 10 μM concentrations of the compounds; verapamil at 10 μM served as a positive control for stimulation of ATPase activity. Significance was determined by a one-way ANOVA with a test for multiple comparisons. Asterisks denote significant difference from the DMSO control, where p<0.05 (*), or p<0.01 (**). ns; not significant.

### Ferroptosis Inducers Inhibit P-gp- and ABCG2-Mediated Transport

As targeted therapies are known to act as inhibitors of ABC transporters, we next characterized the ability of the FINs to act as inhibitors of P-gp or ABCG2. At a concentration of 10 μM, erastin2, ML-162, GPX4 inhibitor 26a, and PACMA31 inhibited P-gp-mediated rhodamine 123 transport, resulting in a 5- to 10-fold increase in rhodamine fluorescence in MDR-19 cells (Fig. 5A). In ABCG2-expressing R-5 cells, GPX inhibitor 26a had the greatest effect on purpurin-18 efflux, while piperazine erastin, ML-162, PACMA31 and RSL3 also significantly inhibited purpurin-18 efflux (Fig. 5B). These results suggest that drug-drug interactions might occur during treatment with FINs.

**Fig. 5.**
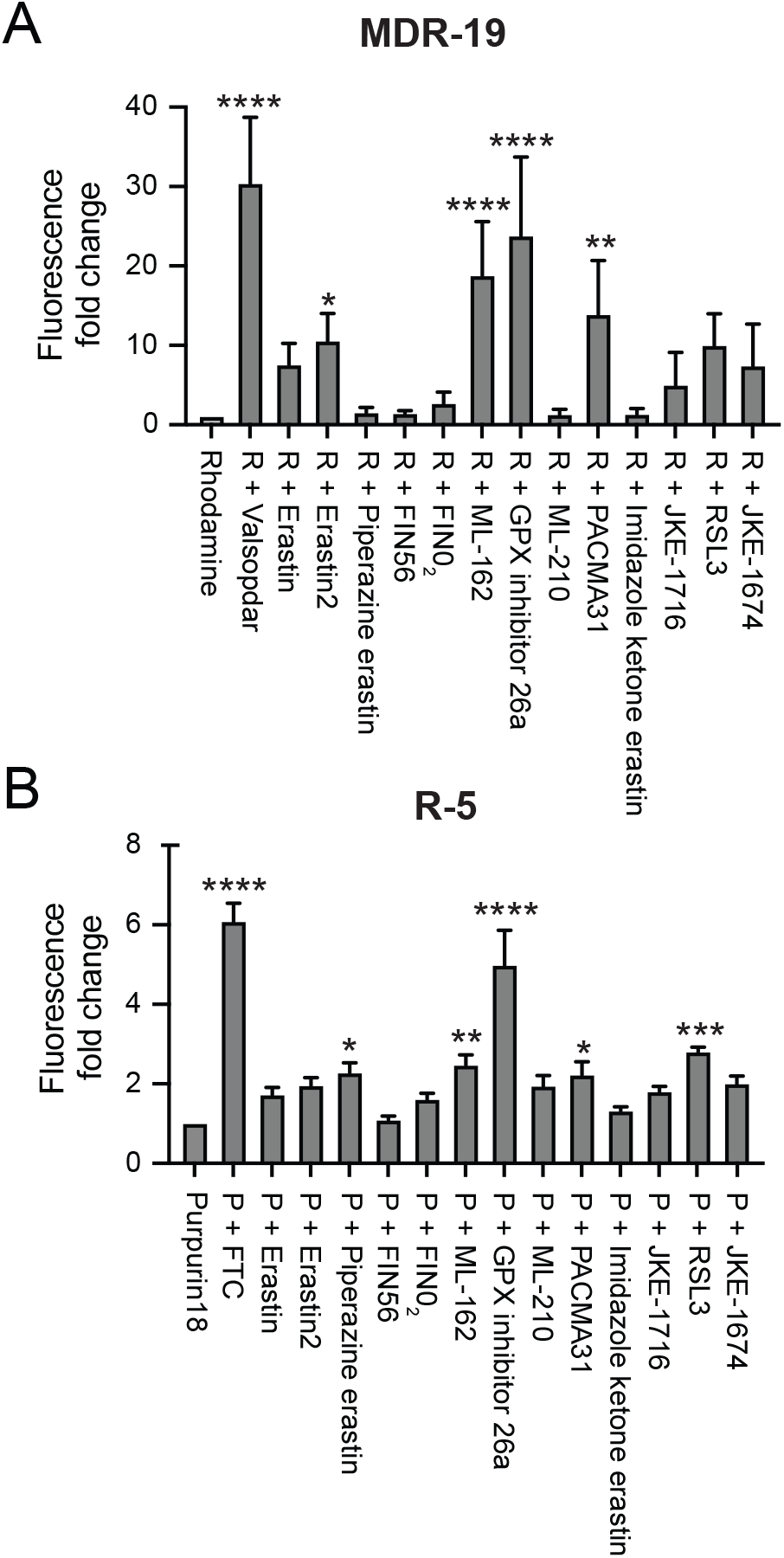
FINs inhibit P-gp and ABCG2 transport activity. P-gp-overexpressing MDR-19 cells (A) or ABCG2-overexpressing R-5 cells (B) were incubated with 0.5 μg/ml rhodamine 123 or 15 μM purpurin-18, respectively, in the absence or presence of specific inhibitor (10 μM valspodar for P-gp and 10 μM FTC for ABCG2) or 10 μM concentrations of the FINs for 30 min after which the medium was removed and replaced with substrate-free medium with or without the inhibitor. Cells were then incubated for an additional 1 h. Inhibition of P-gp or ABCG2 was determined by calculating the fold increase in intracellular fluorescence, with fluorescence levels in cells incubated with rhodamine or pupurin-18 alone assigned a value of 1. Significance was determined from three independent experiments using a one-way ANOVA with a test for multiple comparisons. Asterisks denote significant difference from the rhodamine or purpurin-18 control, where p<0.05 (*), p<0.01 (**), p<0.001(***), or p<0.0001(****). R; rhodamine, P; purpurin-18.

## DISCUSSION

Ferroptosis induction by small molecules is a novel way to induce cell death in cancer cells. Despite the recent proliferation of papers describing novel molecules that can induce ferroptosis (Gaschler et al., 2018; Chen et al., 2020; Xu et al., 2021; Wang et al., 2023), very few studies have addressed potential interactions with ABC transporters that might limit bioavailability or brain penetration. In our study, we found that the FINs imidazole ketone erastin and piperazine erastin are transported by P-gp, suggesting that their oral bioavailability and/or brain penetration may be compromised. Additionally, we found that the FINs ML-162, GPX inhibitor 26a, and PACMA31 act as inhibitors of P-gp. Interestingly, the most potent P-gp inhibitor, GPX inhibitor 26a, was also the most potent ABCG2 inhibitor, suggesting that treatment with FINs may cause drug-drug interactions.

Our findings regarding erastin differ from those of Zhou et al. who reported erastin as a P-gp substrate (Zhou et al., 2019). However, we note that they used a P-gp-expressing cell line that was generated by gradually increasing exposure to paclitaxel (Zhou et al., 2019). While selection with paclitaxel can lead to P-gp overexpression, other mechanisms of resistance can arise (Orr et al., 2003). It cannot be ruled out that other mechanisms besides efflux by P-gp may have caused the resistance to erastin observed by Zhou and colleagues. Unfortunately, they did not perform cytotoxicity assays in the presence of a P-gp inhibitor, which would have confirmed the role of P-gp. In contrast, we did not observe erastin resistance in cells that were transfected to express P-gp without selection with an anticancer drug, and this result was confirmed in a selected cell line. We thus conclude that erastin is not a P-gp substrate and that the cell line used by Zhou et al. may have another mechanism at work that can confer resistance to erastin.

It is not surprising that the FINs, which are essentially targeted therapies, interact with drug transporters, as many targeted therapies have been shown to either be substrates or inhibitors of P-gp or ABCG2 (Shukla et al., 2012). The BCR-ABL inhibitors imatinib, nilotinib, dasatinib, and bosutinib have been found to be substrates of P-gp and ABCG2 at low concentrations, while they act as inhibitors of the proteins at higher concentrations (Hegedus et al., 2009; Tiwari et al., 2009; Dohse et al., 2010; Eadie et al., 2013). Overexpression of *ABCB1* has been demonstrated in tumor samples obtained from patients whose tumors have developed resistance to the ALK inhibitor ceritinib in the absence of secondary ALK mutations (Katayama et al., 2016). Ceritinib has also been reported to inhibit P-gp- and ABCG2-mediated transport (Hu et al., 2015). Additionally, both P-gp and ABCG2 have been shown to confer resistance to several structurally different aurora kinase inhibitors (Guo et al., 2009; Michaelis et al., 2015).

The ability of P-gp and ABCG2 to affect brain penetration of targeted therapies has most dramatically been demonstrated in mouse models in which the *ABCB1* homologs *Abcb1a* and *Abcb1b* are knocked out, the ABCG2 homolog *Abcg2* is knocked out, or all of the homologous transporters have been deleted. Brain concentrations of the Janus kinase 1/2 inhibitor momelotinib 24 h after oral administration were 6.5-fold, 3-fold and 48-fold higher in mice deficient in Abcg2, Abcb1a/b, or Abcg2;Abcb1a/b, respectively, compared to control mice (Durmus et al., 2013). Similarly, 24 h after oral administration of the BCR-ABL inhibitor ponatinib, brain concentrations were 2.2-fold, 1.9-fold and 25.5-fold higher in mice deficient in Abcg2, Abcb1a/b, or Abcg2;Abcb1a/b, respectively, compared to wild-type controls (Kort et al., 2017). Thus, P-gp and ABCG2 can have a profound effect of limiting brain penetration of targeted therapies that are substrates of the transporters. Brain accumulation or oral bioavailability of FIN56, imidazole ketone erastin or piperazine erastin could similarly be affected, as they were found to be transported by P-gp.

In conclusion, we have demonstrated that the FINs FIN56, imidazole ketone erastin, and piperazine erastin are substrates of P-gp, suggesting a potential reduction in oral bioavailability and brain penetration. ML-162, GPX inhibitor 26a and PACMA31 were found to inhibit the transport activity of P-gp and/or ABCG2, suggesting potential drug-drug interactions. Thus, our findings are valuable as these compounds are pursued clinically.

## ACKNOWLEDGEMENTS

DNA cloning support was provided by Vanessa Wall and Carissa Grose of the Protein Expression Laboratory at the Frederick National Laboratory for Cancer Research. We thank George Leiman for editorial assistance. The content of this publication does not necessarily reflect the views or policies of the Department of Health and Human Services, nor does mention of trade names, commercial products, or organizations imply endorsement by the U.S. Government.

## DATA AVAILABILITY STATEMENT

The authors declare that all the data supporting the findings of this study are available within the paper and its Supplemental Data.

## AUTHORSHIP CONTRIBUTIONS

*Participated in research design:* Robey, Ambudkar, Gottesman.

*Conducted experiments:* Frye, Huff, González Dalmasy, Salazar, Carter, Gensler, Robey.

*Contributed new reagents or analytic tools:* Esposito.

*Performed data analysis:* Frye, Huff, González Dalmasy, Salazar, Carter, Gensler, Robey.

*Wrote or contributed to the writing of the manuscript:* Frye, Huff, Salazar, Robey, Ambudkar, Gottesman.

## Footnote

This research was funded by the Intramural Research Program of the National Institutes of Health, the National Cancer Institute.

