## Supplementary material for "The Multidrug Resistance Transporter P-glycoprotein Confers Resistance to Ferroptosis Inducers": Table S

**Table S1.** Cross resistance profile of OVCAR8 and NCI/ADR-RES cells with ferroptosis inducers

| **Compound (µM)** | **OVCAR8 $GI_{50}$** | **NCI/ADR-RES $GI_{50}$** | **OVCAR8+ verapamil $GI_{50}$** | **NCI/ADR-RES+ verapamil $GI_{50}$** |
|---|---|---|---|---|
| Erastin | 2.19 ± 1.66 | 2.07 ± 1.69 | | |
| Erastin2 | 0.15 ± 0.10 | 0.14 ± 0.10 | | |
| Imidazole ketone erastin | 0.36 ± 0.23 | 137.8 ± 45.38 | 0.36 ± 0.23 | 0.55 ± 0.07 |
| Piperazine erastin | 2.15 ± 0.18 | 34.52 ± 9.98 | 2.29 ± 0.05 | 4.11 ± 0.97 |
| FIN56 | 0.47 ± 0.28 | 82.41 ± 4.78 | 0.84 ± 0.51 | 0.28 ± 0.22 |
| JKE-1674 | 1.69 ± 0.65 | 0.81 ± 0.40 | | |
| RSL3 | 0.05 ± 0.04 | 0.06 ± 0.03 | | |
| Fin0$_2$ | 1.79 ± 0.35 | 0.92 ± 0.23 | | |
| PACMA31 | 0.13 ± 0.04 | 0.14 ± 0.08 | | |
| ML-210 | 0.2 ± 0.10 | 0.08 ± 0.04 | | |
| GPX4 Inhib 26a | 0.18 ± 0.13 | 0.14 ± 0.08 | | |
| ML-162 | 0.2 ± 0.03 | 0.16 ± 0.09 | | |
| JKE-1716 | 14.24 ± 4.55 | 8.42 ± 5.00 | | |
| Erastin | 2.19 ± 1.66 | 2.07 ± 1.69 | | |
| Erastin2 | 0.15 ± 0.10 | 0.14 ± 0.10 | | |

Results presented are mean $GI_{50}$ values +/- SEM. Three independent experiments were performed.

Table S2. Cross resistance profile of UO-31 cells and ABCB1-knockout clones

| **Compound** | **UO-31** | **UO-31 B11** | **UO-31** |
| --- | --- | --- | --- |
| Romidepsin (ng/ml) | 42.005 ± 8.53 | 2.801 ± 1.15 | 2.93 ± 0.95 |
| Erastin (µM) | 0.428 ± 0.20 | 0.582 ± 0.02 | 0.582 ± 0.02 |
| Imidazole ketone erastin (µM) | 2.235 ± 0.14 | 0.613 ± 0.05 | 0.583 ± 0.01 |
| Piperazine erastin (µM) | 6.106 ± 0.93 | 2.159 ± 0.31 | 1.949 ± 0.14 |

Results presented are mean $GI_{50}$ values +/- SEM. Three independent experiments were performed.
